# The development of scientific consensus: analyzing conflict and concordance among Avian phylogenies

**DOI:** 10.1101/123034

**Authors:** Joseph W. Brown, Ning Wang, Stephen A. Smith

## Abstract

Recent developments in phylogenetic methods and data acquisition have allowed for the construction of large and comprehensive phylogenetic relationships. Published phylogenies represent an enormous resource that not only facilitate the resolution of questions related to comparative biology, but also provide a resource on which to gauge the development of consensus across the tree of life. From the Open Tree of Life, we gathered 290 avian phylogenies representing all major groups that have been published over the last few decades and analyzed how concordance and conflict develop among these trees through time. Nine large scale backbone trees (including a new synthetic tree from this study) were used for the consensus assessment. We found that conflicts were over-represented both along the backbone (higher-level neoavian relationships) and within the oscine Passeriformes. Importantly, although we have made major strides in our knowledge of major clades, recent published comprehensive trees, as well as trees of individual clades, continue to contribute significantly to the resolution of clades in the avian phylogeny. These findings are somewhat unexpected, given that birds constitute a relatively well-studied and small clade of the tree of life (i.e., Aves). Therefore, our analysis highlights that much work is still needed before we can confidently resolve the less well studied areas of the tree of life.

## Introduction

Large and comprehensive phylogenies (i.e., including hundreds of taxa and based on genome-scale datasets) have become more common as inference methods and sequencing techniques capable of constructing enormous datasets have been developed (e.g., Smith and Donoghue 2008; Rabosky et al. 2013; Zanne et al. 2014; Prum et al., 2015; Simion et al., 2017). These phylogenies have, in many cases, given fresh views to macroevolution and transformed our ability to address diverse sets of comparative biological questions from diversification, morphological evolution, and rate heterogeneity (Brockington et al., 2015; Lin et al., 2016; Scholl and Wiens, 2016). Comprehensive phylogenies that include all or nearly all taxa constructed from supertree techniques also provide a means of determining where data collection efforts should be focused (Davis and Page, 2014; Jetz et al., 2012; Hinchliff et al., 2015). While these trees may facilitate interesting biological inquiries, they also provide a resource by which we can better understand the development of consensus in the community through data acquisition and comparison (e.g., Davis and Page, 2014). There have been efforts to better understand the development of conflict and concordance among trees, with examination having been conducted primarily with molecular data (e.g., Hinchliff and Smith 2014; Smith and Stamatakis 2013; Smith et al. 2015). Nevertheless, phylogenetic resources, including TreeBASE (Sanderson et al., 1994) and more recently the Open Tree of Life (Hinchliff et al., 2015; McTavish et al., 2015), are now available to better analyze how inferred relationships across the tree of life have changed through time.

The Open Tree of Life project has provided the community with several important resources. The Open Tree Taxonomy (hereafter OTT; Rees and Cranston, 2017), unlike many others available, attempts to include only phylogenetically appropriate taxa (e.g., without *incertae sedis*). It is also more comprehensive than other more commonly used taxonomies (e.g., NCBI) as it includes taxa regardless of whether they have molecular data associated. The Open Tree of Life also constructs and serves a draft tree of all described species (Hinchliff et al., 2015), the source data of which consists of published source trees identified, uploaded, and edited by the community. This resource, while continually improving, provides significant opportunities to address broad evolutionary questions that previously would have been impossible. Finally, the project also, openly, provides the database of published phylogenies that have been curated by the community (McTavish et al., 2015). Importantly, the taxa included in each phylogeny have been mapped to a common taxonomy (i.e., OTT), which allows for comparisons to be performed across datasets without an additional tedious and error prone step of name reconciliation. Instead, this reconciliation has already been performed by those who uploaded the tree, often researchers with close knowledge of the focal organisms.

Here, by utilizing the tree database from the Open Tree of Life, we assess the concordance and conflict among the growing number avian phylogenies that have been published during the last few decades. Methods that are used in this study can also be applied to other living groups on Earth based on the Open Tree of Life resources. As the most diverse extant tetrapod lineage with ∼10,800 recognized extant species (Gill and Donsker, 2016) [and potentially more than twice as many cryptic lineages; Barrowclough et al. (2016)], birds have experienced a rapid radiation at deep clades where extremely short internodes exist (Hackett et al., 2008; McCormack et al., 2013; Burleigh et al., 2015; Suh, 2016; Reddy et al., 2017). Although substantial progress has been made on reconstruction of the Aves phylogeny, discovering successive divergence of three monophyletic groups [i.e., Palaeognathae (the tinamous and flightless ratites), Galloanserae (game birds and waterfowl), and Neoaves (all other living birds), Groth and Barrowclough, 1999; Cracraft et al., 2004], resolving the avian phylogeny (especially within Neoaves) has continued to prove a difficult task of the evolutionary biology community since the pioneering efforts of Sibley and Ahlquist (1990).

Explicitly assessing progress in avian phylogenetic inference has recently come to the forefront of the community. By constructing a consensus tree based on six genome-scale phylogenies from five independent studies (i.e., Hackett et al., 2008; McCormack et al., 2013; Jarvis et al., 2014; Suh et al., 2015; Prum et al., 2015), Suh (2016) assessed the reproducibility of various avian phylogenetic hypotheses. Due to the overwhelming conflict among the source trees used (i.e., no higher-level clade could be supported by at least two out of the six trees), Suh (2016) suggested that the very onset of the neoavian radiation produced an irresolvable nine-taxon hard polytomy. Reddy et al. (2017) constructed a nearly identical summary consensus tree to Suh (2016) using a smaller sample of three major hypotheses (i.e., Jarvis et al., 2014; Prum et al., 2015; Reddy et al., 2017), but are more optimistic that more realistic biological-modelling and, importantly, careful selection of data type, will enable further progress. We note that none of the trees considered by Suh (2016) or Reddy et al. (2017) had reasonable sampling of Passeriformes (songbirds; roughly 60% of extant avian species), so potential conflict could not be ascertained within that clade. To date, these and other studies have mainly focused on identifying causes of conflict, attributing tree differences to various factors including incomplete lineage sorting (ILS; Jarvis et al., 2014; Suh et al., 2015), differences in data type (Jarvis et al., 2014; Reddy et al., 2017), and insufficiency of taxon sampling (Prum et al., 2015). However, while these issues of inference are important to keep in mind for future research, little effort has been made to summarize the development and growth of consensus through time when considering the corpus of published phylogenetic hypotheses.

In this study, eight large-scale avian trees [i.e., Sibley and Ahlquist, 1990 (DNA-DNA hybridization); Livezey and Zusi, 2007 (morphology); Hackett et al., 2008 (nuclear DNA); Jetz et al., 2012 (GenBank mining); Davis and Page, 2014 (supertree); Jarvis, et al., 2014 (genomes); Hinchliff et al., 2015 (synthetic tree); Prum et al., 2015 (phylogenomic)] published in different time intervals are used as valuable backbones to assess trends of consensus and conflict. Additionally, after filtering 290 source trees from the Open Tree of Life, we constructed a new comprehensive synthetic bird tree based on certain tree choice criteria (by taxon group, sampling scale, and year of publication) and use it for assessment as the largest avian tree to date. This synthetic tree also serves as a resource for other researchers, and as a point from which we can compare future comprehensive avian phylogenies.

## Methods

### Source trees

Avian phylogenetic hypotheses that have been published in the last few decades were curated through the Open Tree of Life online curator (https://tree.opentreeoflife.org/curator), following the protocol of Hinchliff et al. (2015). In general, published trees (as newick, NEXUS, or NeXML format) were obtained by appealing to authors, or imported from TreeBASE (Sanderson et al. 1994) and Dryad. We attempted to incorporate the source trees from the Davis and Page (2014) supertree study. However, trees from this resource were found to be some form of consensus (e.g., between parsimony and maximum likelihood) hypothesis and/or included unsampled taxa (both extinct and extant) from the Davis and Page (2014) taxonomy. In sum, these trees reflected neither a specific hypothesis nor the extent of sampling of the original publication, and so were not included here. The full species-level tree (i.e., the “Tapestry” tree) of Sibley and Ahlquist (1990) has, to our knowledge, never been available in electronic format. As part of this study, JWB constructed the tree with branch lengths from Figs. 357-368,371-385 of Sibley and Ahlquist (1990); this tree is now freely available from the Open Tree of Life curator (study: ot_427). The taxon labels for each source tree sampled here were mapped to the Open Tree of Life taxonomy (i.e., OTT) and trees were rooted with outgroup based on the original study.

In total, 290 avian phylogenetic hypotheses were gathered from the existing resources in the Open Tree of Life. They are all available in the git-based phylesystem repository (McTavish et al., 2015; https://github.com/OpenTreeOfLife/phylesystem). The distribution of trees sampled through time (Figure 1) reflects data availability rather than research effort, as historically phylogenetic hypotheses have not been archived in machine-readable formats (Stoltzfus et al., 2012; Drew et al., 2013). Among the sampled trees, seven historically trees were used as backbones for later concordance and conflict analyses: Sibley and Ahlquist (1990), Livezey and Zusi (2007), Hackett et al. (2008), Jetz et al. (2012), Davis and Page (2014), Jarvis et al. (2014), and Prum et al. (2015). It is noteworthy that the Jetz et al. (2012) tree we used is the one with genetic data only (6,670 tips) and constraint on Hackett et al. (2008) backbone. In addition to the seven trees above, the Open Tree of Life synthesis tree version 7 (hereafter Opentree7, updated in Sep 2016; Hinchliff et al., 2015) was also included as one of the backbone resources.

**Figure 1.**
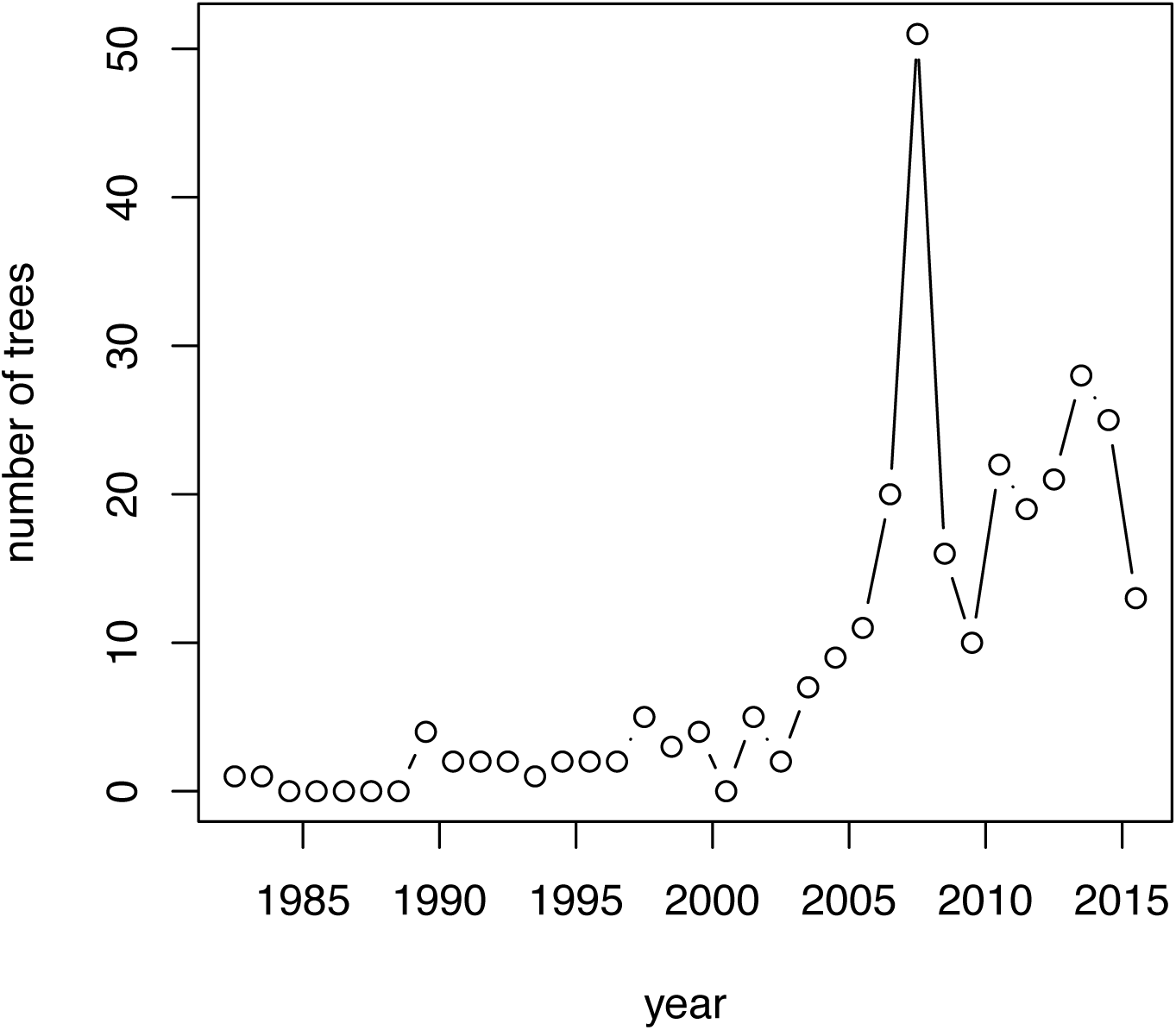
The number of published avian trees per year available in the Open Tree of Life tree repository.

### Construction of a new synthetic tree of Aves

In addition to the individual phylogenetic trees that we collected from the Open Tree of Life, we also assembled a novel synthetic avian tree using the “propinquity” pipeline from Redelings and Holder (2017). This supertree method takes as input a taxonomy tree (i.e., OTT) and a set of ranked source trees. Of the 290 avian trees collected above, 183 were selected as reflecting community consensus about phylogenetic hypotheses. In general, the method will construct a supertree that displays the largest number of input tree edges and minimize the amount of information that does not come from input trees. Moreover, by ranking input trees by certain criteria, a group can be recovered from higher-ranked trees when conflicts exhibit in the source trees. Because synthesis relies upon supertree construction, we excluded both superseded trees (i.e., trees that have been proven to be incorrect by subsequent studies) and previous supertrees. After grouping the bird resource trees into separate focal clades (e.g., by order), we ranked the 183 bird trees based on mixed criteria, such as date of publication, extent of taxon and character sampling, and degree of taxonomic overlap. The set of trees used for the construction of the new synthesis are listed in Supplementary Information. As to conflicting clades, JWB and NW performed the tree ranking order to make sure confident groupings were put in higher ranks. The taxonomic tree with all taxa, derived from OTT, was then used to maximize leaf set for comparison. After dividing the full data set into sub-problems based on uncontested taxa (that is, taxa from OTT that are not conflicted by any source tree), propinquity constructs a summary tree for each subproblem and grafts all summary trees into a single supertree. This result was then grafted with taxonomy-only taxa to produce a complete synthetic tree. See Redelings and Holder (2017) for more details.

### Conflict and concordance analyses

In total, nine major phylogenies (eight published and one new synthetic tree as described above) were used to conduct conflict and concordance analyses. To compare these trees with the 290 source trees, we added the comprehensive OTT set of taxa to each tree by using the same synthesis approach (see above) but with only the taxonomy and the one phylogeny in question. Adding the full OTT set of taxa to the focal trees ensures an overlap of tip sets with each of the 290 source trees. The result was a comprehensive tree with only resolution of the single tree and the taxonomy (i.e., the synthesis procedure preserves all the inferred relationships of the original publication). Conflict and concordance analyses were conducted using the corresponding tools utilized by the Open Tree of Life web service (available from https://github.com/OpenTreeOfLife/reference-taxonomy). Python scripts to perform the analyses are available in the bitbucket repository (https://bitbucket.org/blackrim/opentree_birds). For these analyses, concordant and conflicting edges were identified as in Smith et al. (2013; 2015) and Redelings and Holder (2017). More explicitly, because all sources trees are rooted, a source tree edge *j* defines a rooted bipartition *S*(*j*) = *S*_in_ | *S*_out_, where *S*_in_ and *S*_out_ represent the tip sets of the ingroup and outgroup, respectively. For a given edge in tree *A*, concordance/conflict with source tree *B* thus involves the overlap of tip sets. We define concordance between *A* and *B* (′*A displays B′* in Redelings and Holder (2017)) when *B*_in_ ⊂ *A*_in_ and *B*_out_⊂ *A*_out_ (that is, ingroup tip sets overlap, and outgroup tip sets overlap). [We note that Redelings and Holder (2017) use ⊆ rather than ⊂ because they deal with the more general case where individual trees may have incomplete tip sets. Because we synthesize OTT with each source tree (above), we ensure that each tree has a complete tip set]. On the other hand, edges in trees *A* and *B* are identified as conflicting if none of the following are empty: *A*_in_ ∩ *B*_in_, *A*_in_ ∩ *B*_out_, or *B*_in_ ∩ *A*_out_ (that is, there is reciprocal overlap in the ingroup and outgroup across trees; we note that Redelings and Holder (2017) have a typo in this definition). We computed edge-specific values of concordance and conflict for each of the nine major trees against all 290 source trees. All analyses were conducted on the focal trees with comprehensive taxa added. Finally, we summarized the results on the eight original published tree topologies (i.e., removing the comprehensive taxa that were not included in the original analysis) and the new synthetic tree.

## Results and Discussion

### The synthetic tree of Aves

The synthetic tree constructed in the current study contains 13,579 tips and 10,795 internal nodes, leaving 2,782 nodes (13,577-10,795, assuming a fully binary tree) to be resolved by future studies. We note that this tip count is higher than the ∼10,800 species recognized by Gill and Donsker (2016). OTT is a synthetic taxonomy comprised of numerous source taxonomies that sometimes disagree on the taxonomic status (e.g., species vs. subspecies) or name of a taxon. As a result, there are duplicated taxa in OTT, although this is not expected to influence our results. Comparatively, Opentree7 had more tips (13,756; due to a different version of OTT) but far fewer internal nodes (7,157). Although the changes in taxonomy since Opentree7 have caused the loss of tips in our synthetic tree (probably due to improvements in name reconciliation in OTT), our new synthetic tree resolved ∼ 3,500 more nodes than Opentree7 through inclusion of more source trees (183 vs. 77). The higher-level inter-ordinal relationships shown by the synthetic tree follow mostly that of Prum et al. (2015), which possessed the highest source tree rank regarding the backbone topology. Other studies focusing on individual clades (i.e., order- or family-specific) help to resolve lower level relationships towards the tips. Thus, our new synthetic tree provides a better resource for researchers to conduct evolutionary analyses on birds, and it also serves as a point on which we can measure future improvements in our knowledge of the evolution of this group. However, the new synthetic tree lacks branch lengths due to the difficulty of incorporating branch length information into the synthesis supertree procedure given that the input trees vary in branch length presence, data type (i.e., DNA vs. morphology), branch length measurement (i.e., number of changes vs. time), and scale. While branch lengths and divergence times are not the focus of this study, future studies should examine new ways to incorporate divergence times to increase the utility of this resource.

### The distribution of phylogenetic conflict

One major aim of this study is to examine the distribution of concordance and conflict across nine major bird phylogenies. Except for that from Sibley and Ahlquist (1990), all the historically important backbone trees we considered were constructed since 2007. Each tree, at the time of publication, represented major improvements in data collection and analysis, and therefore also on the hypotheses about avian evolutionary history. The most recent trees agree on many relationships despite being built using different methods [i.e., one tree is based on morphological data (Livezey and Zusi, 2007), four are built with different scales of molecular data (Hackett et al., 2008; Jetz et al., 2012; Jarvis et al., 2014; Prum et al., 2015), while the rest were based on supertree methods (Davis and Page, 2014; Opentree7; our synthetic tree)]. Given that Sibley and Ahlquist (1990) and Prum et al. (2015) represent the earliest and most recent comprehensive hypotheses, respectively, results from these trees and the new synthesis tree are discussed in more detail for comparison.

There are more conflicting edges (warmer colours, Figure 2) for higher-level relationships (i.e., the backbone from Neognathae to Passeriformes) on the Sibley and Ahlquist (1990) tree. This is to be expected given that many phylogenies have been published since 1990 that have contradicted the Sibley and Ahlquist (1990) “tapestry” hypothesis (see also discussion in Harshman, 1994). For instance, the rooting of this tree, with Galloanserae sister to Paleognathae, has been refuted by all subsequent studies. The new synthetic tree has several higher relationships (i.e., toward the base of Neoaves) that have higher conflict than that of Prum et al. (2015), but this is likely due to the comprehensive nature of the tree. This pattern is largely consistent with the neoavian polytomies posited by Suh (2016) and Reddy et al. (2017), although these studies only considered a small number of phylogenomic studies for comparison. Many of the conflicts among deepest neoavian relationships are also shared by Jetz et al. (2012), which contributes many of the edges to the synthetic tree due to the extensive taxonomic sampling (Figure 3). However, the Prum et al. (2015) tree (Figure 2) has comparatively fewer conflicting edges toward the base of Neoaves. This may simply be due to significantly lower taxon sampling (i.e., fewer sampled lineages means a smaller potential number of conflicts).

**Figure 2.**
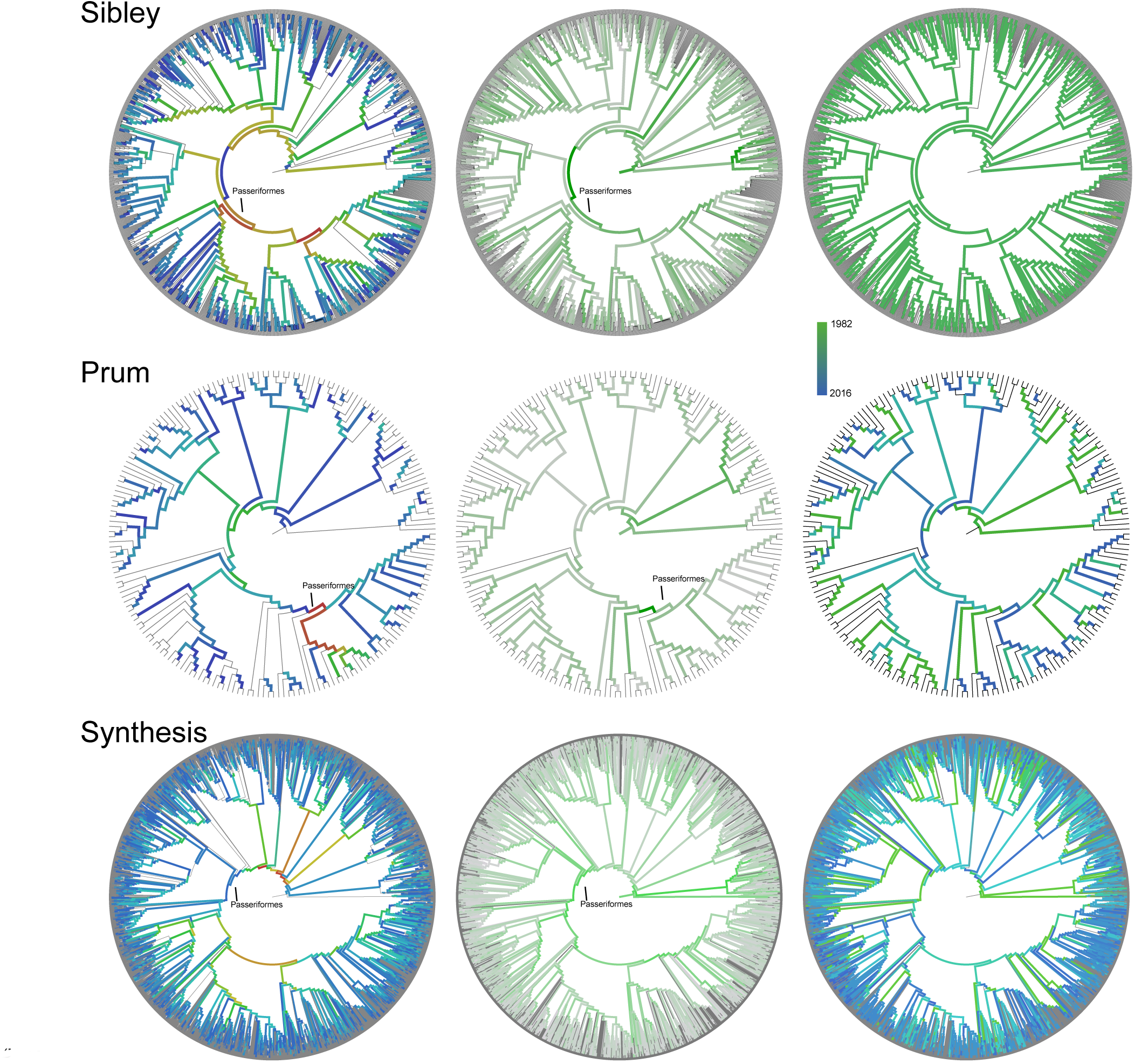
Conflicting (left, warmer colour represents more conflict), concordance (center, darker green represents more concordance), and chronological contribution (right) analyses based on three backbone trees. Conflict and concordance are measured as the number of source tree edges that conflict/support a given focal tree edge. Chronological contribution is measured as the first year in which the focal edge appeared in a published tree.

**Figure 3.**
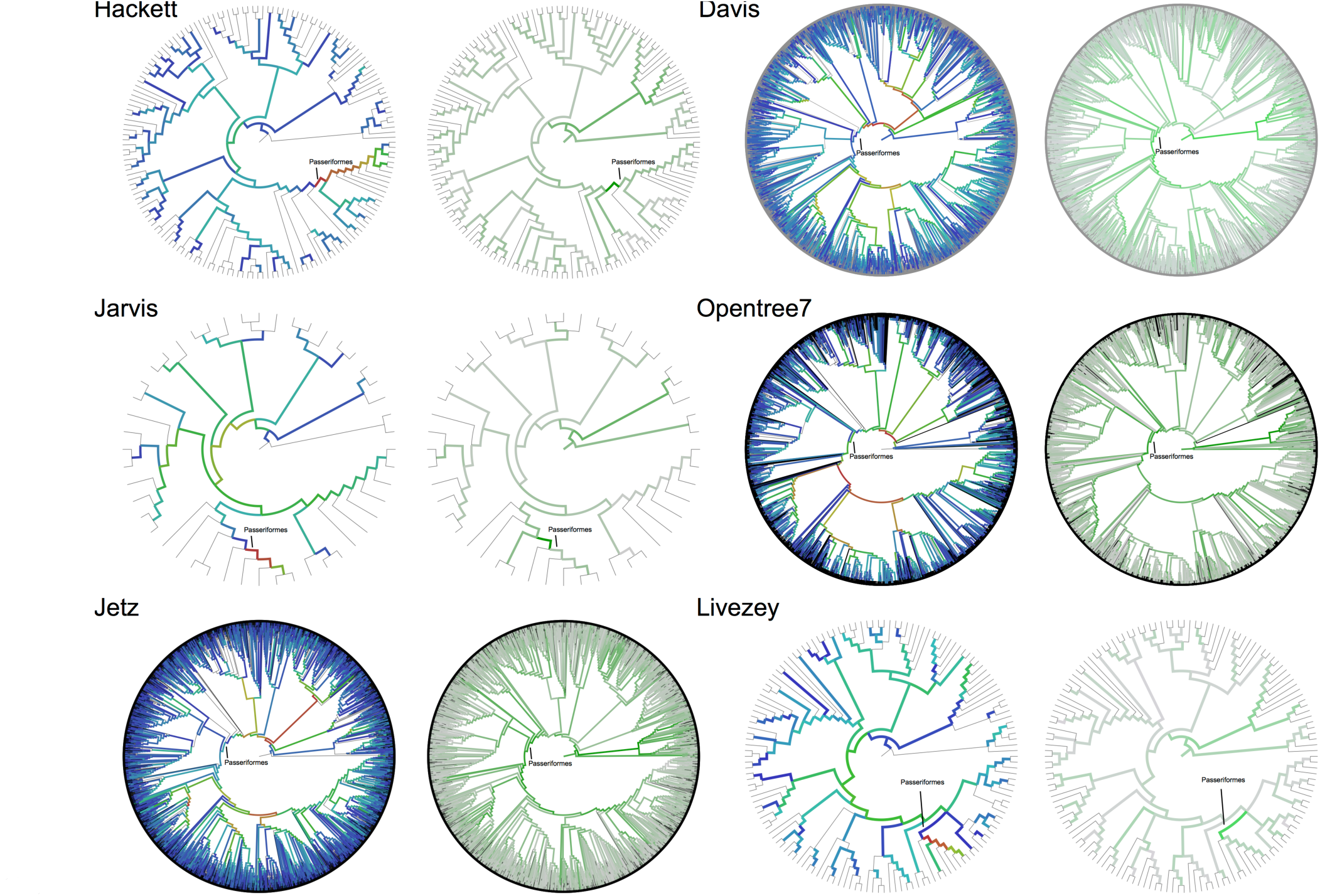
Distributions of conflict and concordance across the six-remaining major avian phylogenetic hypotheses. See Figure 2 for description of colours.

A more general pattern of conflict across the focal trees involved the Passeriformes (songbirds). In fact, although the monophyly of Passeriformes is largely supported, the relationships that diverge soon after crown Passeriformes (i.e., within the oscines) are highly contested among the source trees. Sibley and Ahlquist (1990) have the highest conflict among edges surrounding the Passeriformes clade, with six having 29-37 conflicting trees and 13-30 supporting trees. Similar patterns with strong conflict surrounding the edges of Passeriformes were exhibited on the other focal trees. Prum et al. (2015) showed strong conflict at the base of Passeriformes, with exceptionally high conflict leading to oscine songbirds. The synthetic tree showed fewer conflicts at the base of Passeriformes, implying that many of the source trees that contributed to the synthetic tree construction contradict the oscines relationships in Prum et al. (2015), possibly due to the limited taxonomic sampling in the latter. We note that strong phylogenetic conflict within Passeriformes was not evident in the studies of Suh (2016) and Reddy et al. (2017), as they only considered trees focused on inter-ordinal relationships. Although conflict and concordance within Passeriformes has not previously been explicitly evaluated, individual assessments of phylogenetic uncertainty have hinted that this may involve a number of problematic relationships. For example, Burleigh et al. (2015) report that for clades with < 50% bootstrap support, almost half of the families (195/399) and two thirds of the genera (32/47) were within the oscine songbird clade. Given that Passeriformes represents ∼60% of extant avian species, these results together suggest that there is much work to be done in avian phylogenetics than resolving the neoavian backbone.

### The resolution of the Aves phylogeny through time

The trees on which we conduct conflict and support analyses represent either major contributions or comprehensive analyses of the Aves clade. One central question regarding these trees is how does our knowledge about clades change over time. For example, did recent genomic studies such as Prum et al. (2015) contribute many unique clades to the literature, or were these resolutions simply recapitulating results that had been published previously? Although we are limited by the database of 223 available trees, we can begin to address these questions. When compared to earlier studies, Sibley and Ahlquist (1990) contributes almost all new relationships for bird phylogeny except for three edges (published before 1990) that involve relationships of flycatchers, *Phrygilus*, and *Junco* (Figure 4). It is likely that more edges on Sibley and Ahlquist (1990) had already been published in the older literature. However, we do not have access to many of those trees because they are not available in electronic format. By comparison, the Prum et al. (2015) study provides an updated view of the deep relationships of birds (Figure 4). There are a range of dates (i.e., previous published hypotheses) supporting edges of the Prum et al. (2015) tree. While 143 edges had been reported in previous publications, including many in Sibley and Ahlquist (1990), 47 edges were new to the Prum et al. (2015) tree. This highlights that important contributions are still being made by large-scale genomic and molecular studies.

**Figure 4.**
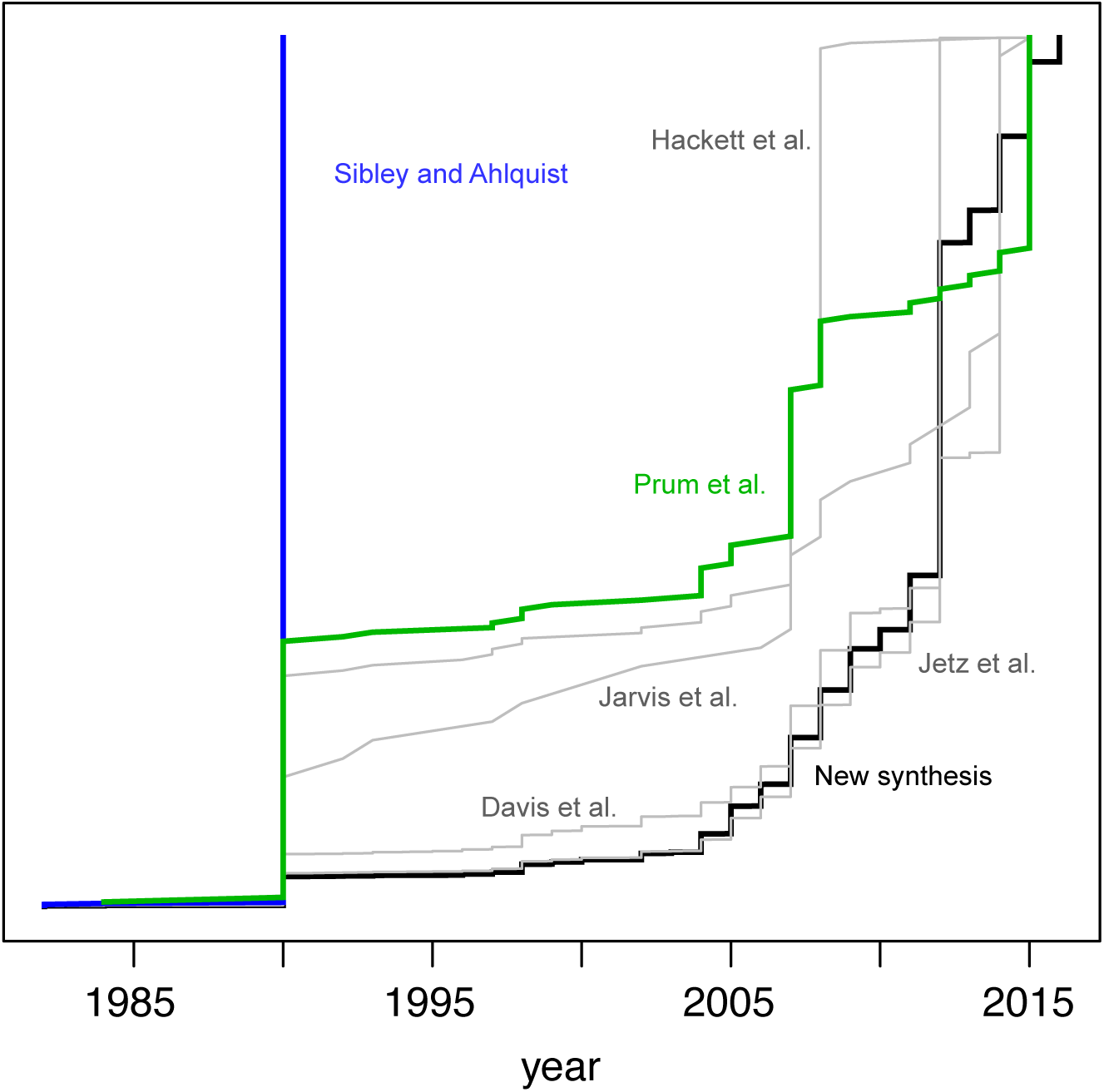
The contribution of source tree support to each of the major avian phylogeny. The x axis represents the source study year that first supports a clade in a focal tree, and y axis is the proportional accumulation of clades on each tree.

The new synthetic tree reported here provides another interesting contrast. Although comprehensive trees have been published before, including Jetz et al. (2012) and Davis and Page (2014), our synthetic tree allows us to address whether new studies are contributing significant new knowledge besides these comprehensive trees. The largest contributing study to the edges of the synthetic tree is Jetz et al. (2012) with 4,820 (Figure 3). However, 1,890 edges were contributed after 2012, and 223 were added from studies published after 2015. This underscores the importance of not only large genomic studies, but also of trees of individual clades to the knowledge of bird phylogeny.

Large phylogenies like those analyzed and created as part of this study have many potential uses. They have been and continue to be used in large scale studies of biodiversity (e.g., Jetz et al., 2012; Cooney et al., 2017). As mentioned above, methods need to be developed to apply dates to the synthetic trees and supertrees that are produced as part of comprehensive analyses (Davis and Page, 2014; Hinchliff et al., 2015; Redelings and Holder, 2017). The development of methods for applying divergence times will dramatically improve the utility of comprehensive trees for further comparative studies. In addition to the larger comprehensive trees, the Open Tree of Life tree repository (McTavish et al. 2015), with phylogenetic hypotheses mapped to a common taxonomy, becomes increasingly useful as it continues to grow. As demonstrated here, these trees can be used to gauge how our understanding changes, whether new studies are contributing new edges, and localize the major sources of conflict. Furthermore, the relationships can serve to construct meaningful prior expectations for the resolution of clades across the tree of life (e.g., as topological priors in a Bayesian reconstruction). It is noteworthy that all these analyses depend on the availability of electronic tree files that are crucially important for any publication that posits phylogenetic hypotheses, but archiving such resources has not been common historically (Stoltzfus et al., 2012; Drew et al., 2013). As we continue to improve our view of the tree of life, it will be instructive to examine how consensus builds across major clades.

## Conclusion

A fundamental goal for the field of evolutionary biology and systematics is the resolution and construction of a complete tree of life. The resources for constructing comprehensive trees (e.g., phylogenetic trees, molecular and morphological data sets, and comprehensive taxonomies) are becoming available and are now of the quality that we can not only begin to construct complete trees, but also refine and identify where more work is needed. Here, we demonstrate that, while we have made major strides in our knowledge of some clades, new studies continue to contribute new edges that resolve previously ambiguous relationships. We make this observation on Aves, a relatively well-studied and small clade of the tree of life. Other parts of the tree of life that are likely to have a lower density of phylogenetic information in the form of published phylogenies or molecular data still need significant more work before they may be comprehensively resolved.

## Acknowledgements

We would like to thank the following for comments on previous versions of this manuscript: Ben Winger, Karen Cranston, and members of the Smith lab. We especially thank Ed Braun for a particularly thorough and thoughtful review of a previous draft. We thank Ben Redelings and Mark Holder for discussions on the definitions of conflict and concordance. JWB, SAS, and NW were supported by NSF DEB AVATOL grant #1207915.

## Data availability

All the software and data used for this study are freely available in repositories online. For the source trees and software used in the analyses, please see https://bitbucket.org/blackrim/opentree_birds.

## Author contributions

JWB and NW gathered the input source trees and constructed the synthetic trees. JWB, NW, and SAS wrote the manuscript and conducted the analyses.

